# Large-scale correlations between eight key double strand break related data sets over the whole human genome

**DOI:** 10.1101/581173

**Authors:** Anders Brahme, Maj Hultén, Carin Bengtsson, Andreas Hultgren, Anders Zetterberg

## Abstract

Eight different data sets, covering the whole human genome are compared with regard to their genomic distribution. A close correlation between cytological detected chiasma and MLH1 immunofluorescence sites with the recombination density distribution from the HapMap project was found. Sites with a high probability of chromatid breakage after exposure to low and high ionization density radiations are often located inside common and rare Fragile Sites (FSs) indicating that the common Radiation-Induced Breakpoint sites (RIBs) may be a new kind of more local fragility. Furthermore, Oncogenes and other cancer-related genes are commonly located in regions with an increased probability of rearrangements during genomic recombination, or in regions with high probability of copy number changes, possibly since these processes may be involved in oncogene activation and cancer induction. An increased CpG density is linked to regions of high gene density to secure high fidelity reproduction and survival. To minimize cancer induction these genes are often located in regions of decreased recombination density and/or higher than average CpG density. Interestingly, copy number changes occur predominantly at common RIBs and/or FSs at least for breast cancers with poor prognosis and they decrease weakly but significantly in regions with increasing recombination density and CpG density. It is compelling that all these datasets are influenced by the cells handling of double strand breaks and more generally DNA damage on its genome. In fact, the DNA repair genes are systematically avoiding regions with a high recombination density. This may be a consequence of natural selection, as they need to be intact to accurately handle repairable DNA lesions.

## Introduction

The Double Strand Break (DSB) repair pathways of human cells are normally very effective. For example, at the common dose during radiation therapy of 2 Gy per treatment, about 75 DSBs are generated in each cell, but on average less than one of these DSBs are lethal, at least in normal tissues. In fact, at this dose level on average less 0.9 % of low ionization density X-ray and electron induced DSBs are lethal for normal cells^1^. Fortunately, due to the genetic instability of most tumor cells, their repair fidelity is significantly lower, so after about 30 radiation treatment fractions a small local tumor (≈10^10^ cells) may be eradicated without severe normal tissue morbidity. It is largely the well functioning Non Homologous End Joining (NHEJ) and not least Homologous Recombination (HR) repair systems that ensure the high fidelity repair of normal tissues. NHEJ is fast but error prone, whereas high fidelity HR, which is mainly active during the late G2-M phase of the cell cycle, may correct mismatches in preparation for mitosis. It is not unlikely that NHEJ, like the meiotic recombination process, also attach a DNA mismatch repair protein, like the MLH1 binding site of meiotic recombination, in order to facilitate and allow efficient proofreading for HR, during the G2-M phase of the cell cycle.

The recombination process is of vital importance for correct segregation of the hereditary material in connection with the crossover of homologues during meiosis leading to the formation of normal haploid gametes. The recombination process is essential to allow the inheritance of new genetic properties and DNA damage and DNA replication often requires the homologous recombination mechanism for high fidelity repair. During chromosome segregation each bivalent pair of homologues needs at least one crossover and the associated DSBs to successfully segregate. The number of exchange events is increasing with chromosomal length and may be as high as five in humans. Recombination events can occur at many sites along a chromosome, but a crossover in one region generally inhibits the formation of another one in close proximity (crossover interference). A repression is also observed in the centromere region. The density of recombination events is not distributed uniformly, but is known to be high at numerous (∼10^5^) recombination “hot spots”, where DSB seem to be preferentially induced^2^. The process behind the recombination events, and the reason why some regions are “hot” while others are not, is largely unknown but may be linked to the need for some genes to be shared as assemblies.

The first method used for analyzing meiotic recombination was cytological analysis of chiasma in human germ cells. With this method Hultén was the first to determine the distribution of chiasma events for individual chromosomes in males^3^ and later also for females. Unfortunately, this approach is limited by the finite optical resolution of light microscopy. The use of immunofluorescence to localize the DNA mismatch repair protein MLH1 is another method for detection of the recombination sites. MLH1 binding sites have been recognized to identify the crossing over locations in mice^4, 5^. The need for correct chromosome segregation and the risk for genomic alterations at the recombination sites may be the reason for the incorporation of a binding site MLH1 in these regions. This method has been used by Barlow and Hultén^6^ in man and later by Sun et al^7^ to produce recombination maps for all male autosomal chromosomes based on the variation in 10 individuals. More recently, Myers et al^8^ used the method of linkage disequilibrium, the non-random association of SNPs (Single Nucleotide Polymorphisms) to produce a fine-scale recombination map in human males. Recombination density data identified by the three different methods were compared to each other for all autosomal chromosomes. The distribution of the recombination sites is clearly nonrandom.

Almost simultaneous with the cytological banding studies by Hultén^3, 6^, a series of papers on the nonrandom distribution of Radiation-Induced Chromosomal Breakpoints (RIBs) of lymphocytes in the G0 and G2 cell cycle phases by Holmberg et al^9–12^. This was seen even if the initial damaging events are known to be largely random over the whole genome, at least with low ionization density radiations whereas densely ionizing particles may generate large clusters of strand breaks along their path. Holmberg et al ^9, 10^ and Kiuru et al^13^ have found that more X-ray RIBs are mapped to G-light bands that are gene-rich transcriptionally more active regions of less condensed chromatin than to G-dark bands.

Fragile Sites (FS) are a common feature of human chromosomes. They can be viewed as non-randomly distributed site-specific loci that are especially prone to forming gaps, breaks or triradial figures in metaphase chromosomes^14^. To date there is approximately 120 known fragile sites, and on the basis of their frequency they are classified into two types, common and rare. “Rare” fragile sites (RFS) are present in less then 2.5% of humans^15^, and they seem not to be implicated in cancer development. In contrast, common fragile sites (CFS) are present on chromosomes of all individuals as a component of normal chromosome structure^15^. They have been known for many years as a chromosomal expression of genetic instability and have been implicated to have a causative role in cancer.

To get an idea how critical or important a specific genomic region really is, it is interesting to study the CpG density in that region. Vertebrate genomes are depleted of the CpG dinucleotide, due to the DNA methylation at the 5-position of cytosine within the CpG dinucleotide. The deamination product of 5-methyl C is T, which easily escapes the repair mechanisms. Over evolutionary time, the C nucleotides in the CpG dinucleotide tend to mutate to T, leaving a genome with CpG deficiency where CpG occur at only one-fifth of its expected mean density^15^. In certain regions of the genome the CpG dinucleotide occur at a density closer to that predicted by local G+C content. These regions or “CpG islands” are often found in promoter regions, or within the first exon of genes^16^. A mutation in an essential region of the DNA should be avoided at all costs, and regions of higher than average CpG density thus are more likely to contain “critical” DNA sequences. The CpG density has a strong association with the recombination rate on the megabase scale, whereas on the kilobase scale they are important but their influence on the recombination rate is diminished^5^. Human DNA copy number alterations have previously been studied mainly by looking at individual genes. With the Representational Oligonucleotide Microarray Analysis^17^ (ROMA) technique it is now possible to scan the entire genome to look for this kind of large-scale variation. ROMA is a suitable method for detecting copy number changes (CNCs) in tumor cells, such as in the presently studied breast cancer patients. Their CNC are large, from the kilobase up to the megabase scale, duplications and deletions sometimes resulting in the transformation of a cell into a cancer cell. The ROMA approach allows a high resolution, on average 35 kb, since the microarrays used presently consist of 83 000 representative probes, covering the whole genome. The technique uses a representation of the genome, acquired by digestion of the DNA with a restriction enzyme, followed by amplification of the fragments by adaptor-mediated PCR. The oligonucleotide probes are designed to be complementary to these fragments and placed in a microarray, where the sample of interest is hybridized against a reference. In this study the data used is from samples acquired from a set of data for breast cancer patients, treated at the Karolinska University Hospital in Stockholm. Recombination events and CNCs have in common that the DNA helix has at some point in time been cut and a segment of DNA has been inserted or deleted.

It would therefore be interesting to correlate the above data sets with the locations of cancer-associated genes, many of which are included in Figure 1. This figure shows the principal genetic pathways, such as those for growth control, DNA damage surveillance and repair, stress response and cell cycle regulation that we know today to be affected in one way or another in practically all cancer patients. In the present study the eight mentioned datasets are compared over the whole human genome to search for relevant correlations and connections.

**Figure 1.**
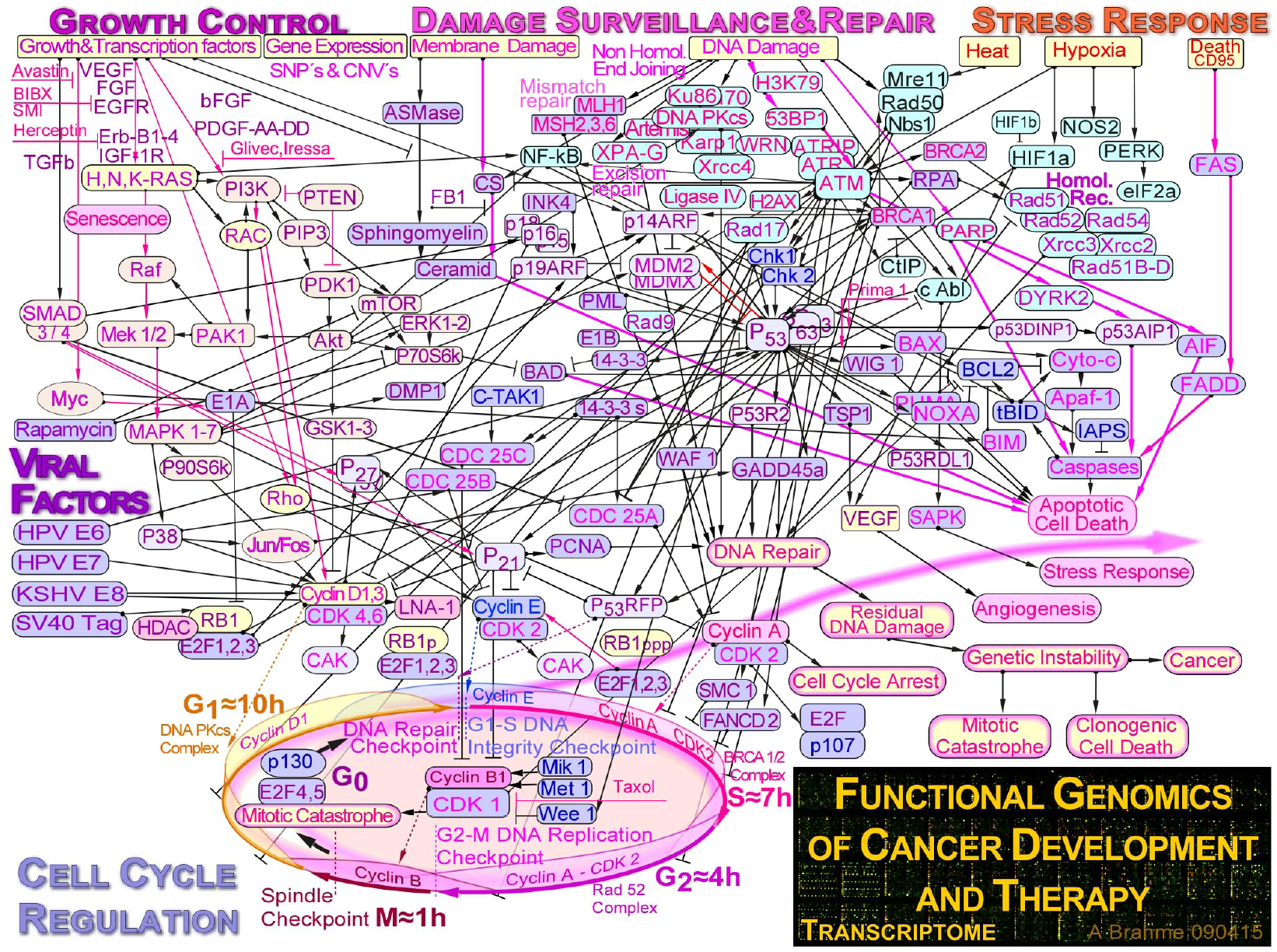
Some key cancer related genes and their complex associated pathways as included in upper lanes of Figure 2–5.

## Material and Methods

### Radiation-Induced Chromatid Breaks (RIBs)

The distribution of X-ray RIBs along individual chromosomes was acquired from Holmberg and Jonasson^9, 10^. Human lymphocytes cultured in vitro were irradiated with X-rays during the G_0_ and G_2_-phase of the cell cycle. The blood samples were obtained from five chromosomally normal individuals (2 males, 3 females). The RIBs were found to be preferentially located in the white R-bands using the banding pattern of the Paris Conference (1971). Later Holmberg and Carrano^10^ found that neutron RIBs in human chromosomes have only partly similar locations as X-ray RIBs. The distribution of RIBs was studied in G_0_-phase lymphocytes irradiated with fast neutrons. With the G-banding technique the breakpoints were mainly located in the pale bands. In the present study the X-ray and neutron RIBs were graphically transferred to an ideogram of the G-banding pattern at the 850 band resolution from NCBI, in June-July 2005 (build 35.1) http://www.ncbi.nlm.nih.gov/mapview/ as shown in the lowest two lanes of Figures 2–5.

**Figure 2.**
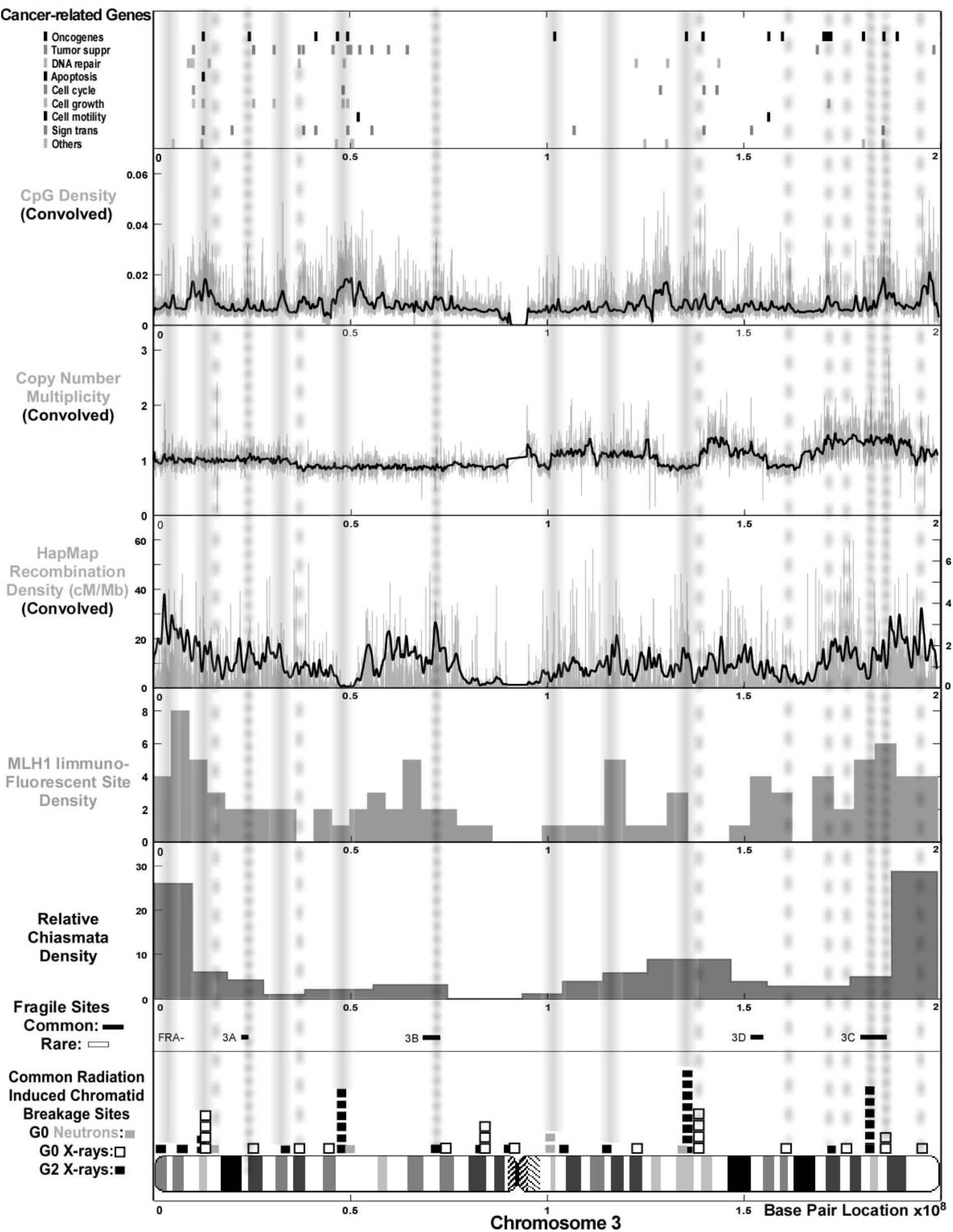
chromosome 3.

**Figure 3.**
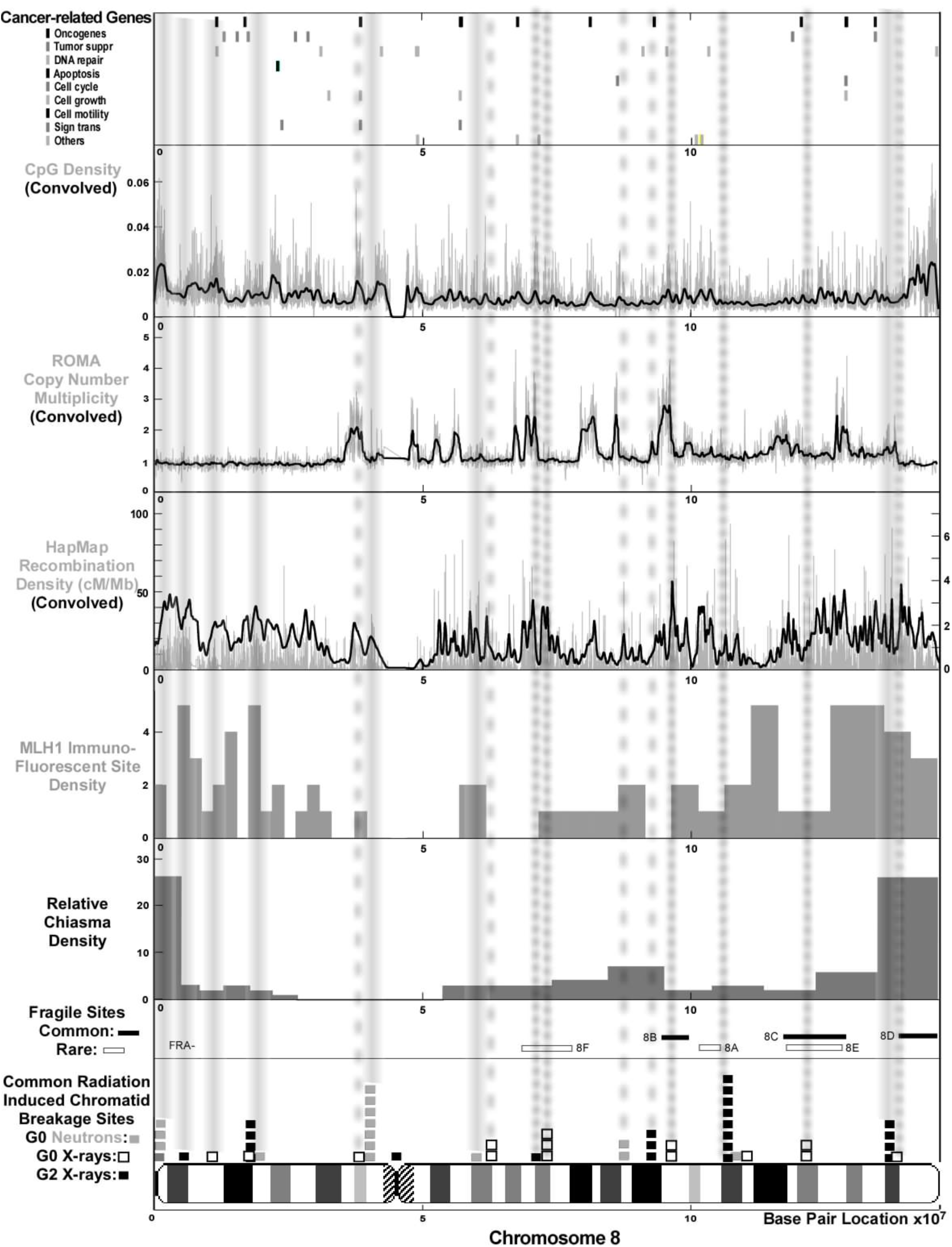
chromosome 8.

**Figure 4.**
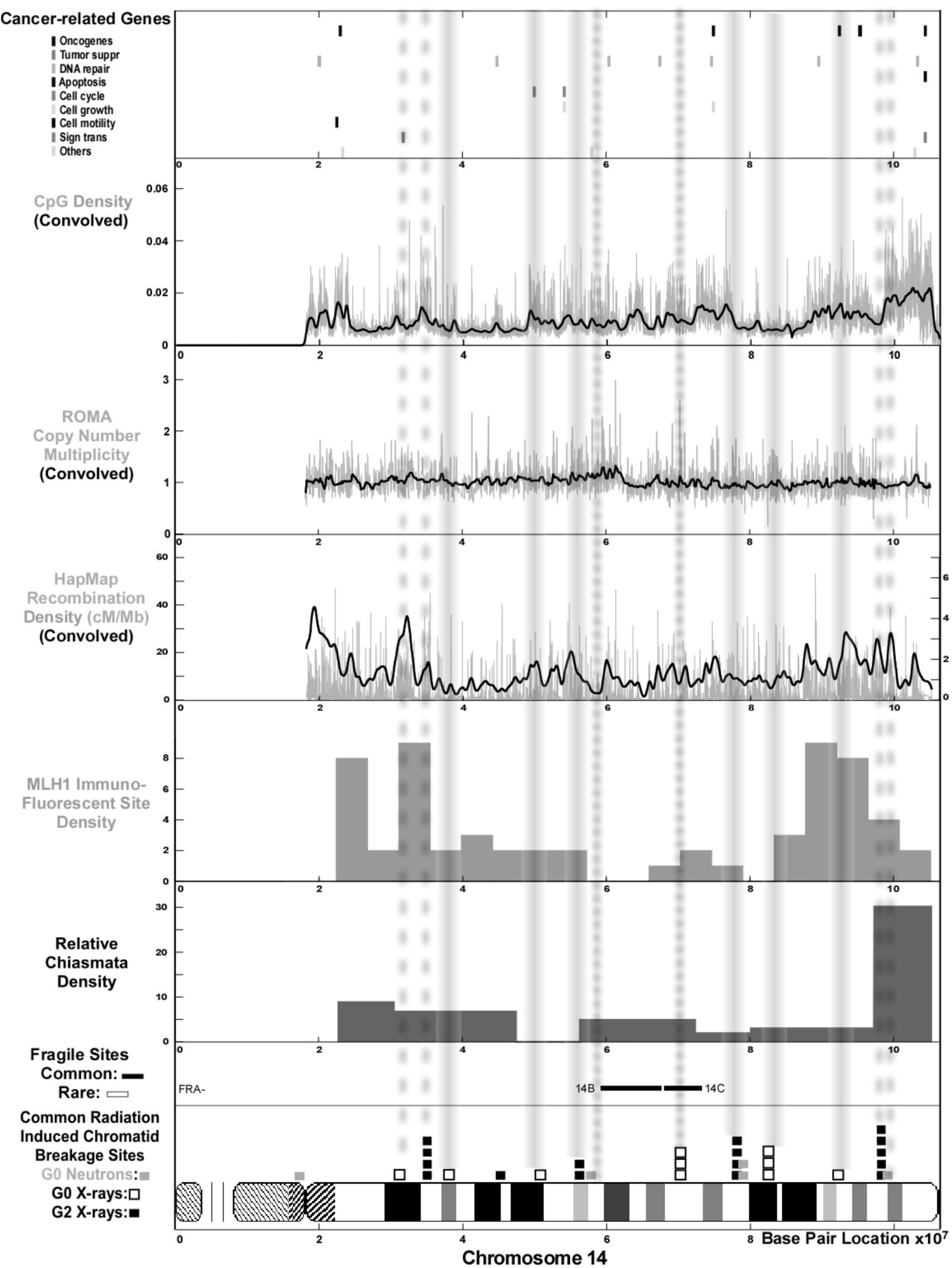
chromosome 14.

**Figure 5.**
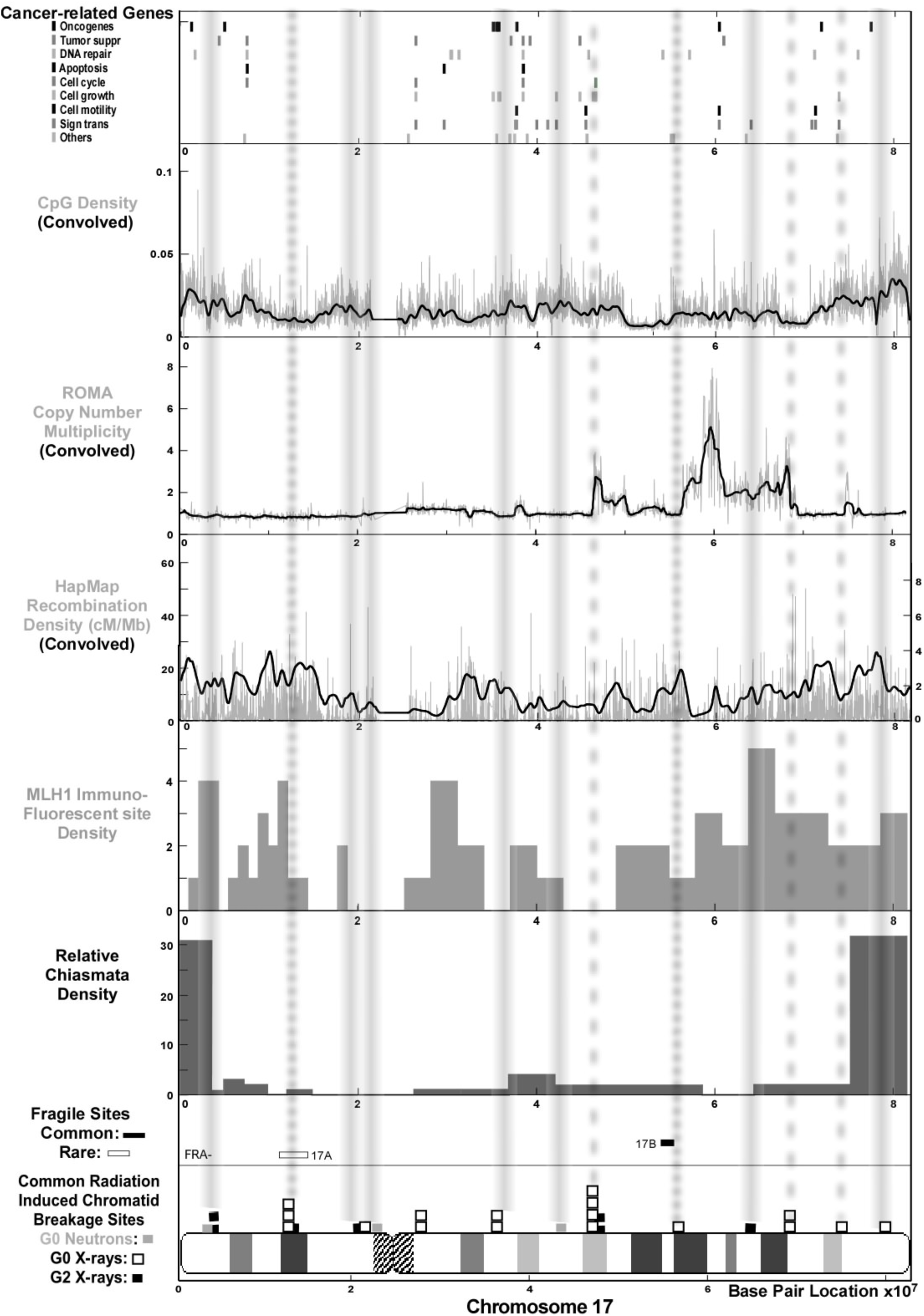
chromosome 17.

**Figure 2–5.** Comparison of common radiation-induced chromatid breakpoints and fragile sites with recombination density distributions, copy number changes, CpG density and cancer associated genes on human chromosomes 3, 8, 14 and 17 respectively. Also included at the bottom of the Figures is the respective G-banding chromosome ideogram at the 850-band resolution and a base pair location scale common for all lanes. It is interesting to see that the radiation induced chromatid breakpoints (RIBs) are commonly located in regions with a high recombination density either as measured through the HapMap project or by MLH1 immunofluorescence or Chiasma Density (Gray Lines, the width of the lines corresponds to the approximat location uncertainty). There is also an interesting correlation, especially on chromosomes 3 and 8, between ROMA detected copy number multiplicity, RIBs and the HapMap recombination density (dashed gray lines, cf also Table 2), as well as between RIBs and C & R FS (cf. FS 3 A, B&C and 8 A, B, C, D&F see the fine dotted gray lines)

**Table 2:**
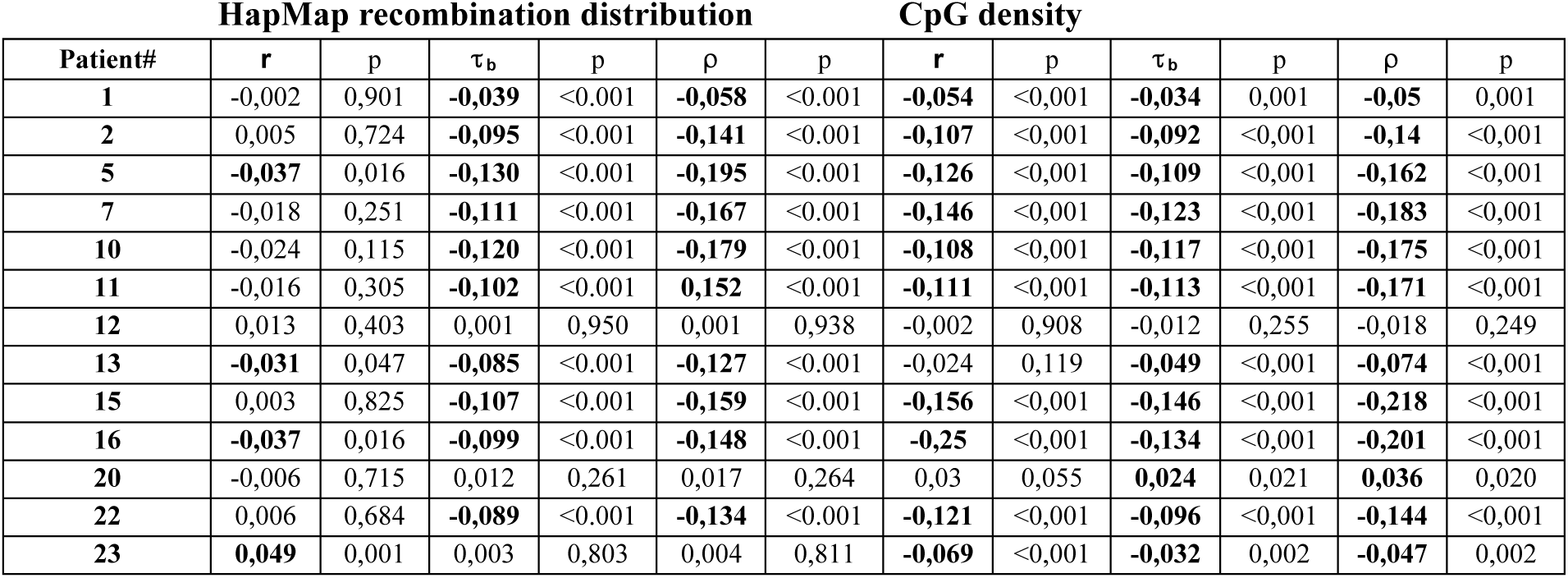
Correlation of HapMap recombination distribution and CpG density with copy number multiplicity in cancer cells on chromosome 8 for 13 individuals with breast carcinoma.

### Fragile Sites (FS)

“Common” fragile sites (CFSs) are also predominantly located in light G bands that are nuclease-sensitive. These regions are characterized by high genomic instability, and chromosome locations of many CFSs coincide with locations of cancer breakpoints and/or cancer-related genes. In a study by Buttel et al^14^ it was found that 67% of the in vitro-induced fragile sites were located in cancer-specific breakpoints, and 72% were co-localized with identified oncogenes. CFS also seem to be hot spots for sister chromatid exchanges^18^. With these properties of the fragile sites, a large-scale comparison with the other structures was performed in this study. An obvious problem is that the fragile sites, like chromosomal breakpoints and chiasma are generally only mapped to a cytological band of a chromosome, making a more detailed correlation analysis complicated. The location of the common and rare fragile sites was taken from NCBI, www.ncbi.nlm.nih.gov/ in June-July 2005 (build 35.1) as shown in Figs 2–5 above the RIBs.

### Recombination, MLH1 immunofluorescence and chiasma distributions

Three different sets of recombination site information were used, based on different methods of detection as shown by the next three lanes of Figure 2–5. Already 1973 Hultén et al^3^ investigated the chiasma distribution at diakinesis in one human male using cells triple-stained by the orcein, Q-and C-techniques. Each chromosome arm was divided into 10 % intervals, in which the chiasma were counted. Later Sun et al^7^ used the immunofluorescence techniques to create a recombination map of the autosomal chromosomes in the human male. The data from Sun et al contains the recombination density in 5% intervals at each arm of the autosomal chromosomes from 100 human male Pachytene stage cells. When estimating finer-scale recombination rates, analysis of linkage disequilibrium (LD)^19^ is the method of choice. Recombination rates based on LD-data and the location of recombination hotspots were downloaded from the HapMap Consortium^20^, hapmap.ncbi.nlm.nih.org. The recombination rate and the hotspots were calculated from the Phase I HapMap data (release 16a) using the methods described in McVean et al^21^ and Winckler et al^22^.

### Copy Number Changes (CNCs)

The Copy Number Changes was acquired from ROMA-data, supplied by Zetterberg, Hicks and Wigler^23, 24^ from tumor samples of 13+12 breast carcinoma patients with poor prognosis. The arrays used for detection had 83 000 probes at an average resolution of 35 kb. The data from Wigler was analyzed with a segmentation algorithm^25^ to help identify the regions of significant CNCs (cf Fig 7). The complete ROMA copy number multiplicity spectrum was in the present study also convolved with a suitable narrow Gaussian filter to reduce random fluctuations and enhance the resolution of small regions of altered multiplicity across the genome. The convolution together with the segmentation algorithm proved to be useful tools for accurately identifying the CNCs. The site for a CNC was defined as the region where the ROMA data had amplitude changes of at least 20% detected either with the segmentation algorithm or the Gaussian filter as seen in the lane above the HapMap data in Figure 2–5. This may correspond to that at least 10% of the cells has a local dual copy number.

### CpG density

The genomic data used to study the local CpG density were downloaded from ftp://hgdownload.cse.ucsc.edu/apache/htdocs/goldenPath/hg17/chromosomes/ which contains the Build 35 of the finished human genome assembly. The positions of the centromeric regions were extracted from the same source. The raw data was then modified as follows: First, an algorithm was written to search through every base. If a G followed a C, the position on the chromosome was given a weight of unity otherwise zero. In this way the CG sequence data was stored in a binary vector. Then each chromosome was divided into equal intervals, with a size of 10 000 bases. The CG density at each interval was defined as the sum of the intervals, divided with the 10 000 bases. The resultant vector was also convolved with a Gaussian filter to get a smoother picture of the variation of the CpG density, as seen in Figures 2–5 above the CNC lane.

### Cancer-related Genes

The list of cancer-related genes used in this study is an assembly from different sources. The oncogenes and tumor suppressor genes and the list of DNA repair genes was attained from: (http://www.binfo.ncku.edu.tw/cgi-bin and http://www.cgal.icnet.uk/DNA_Repair_Genes.html now updated at http://sciencepark.mdanderson.org/labs/wood/dna_repair_genes.html). Other genes such as genes involved in apoptosis, cell-cycle control, growth control, cell motility and signal transduction, is the result of a mapping of pathways for genes, involved in cancer development and therapy, as shown in Figure 1. The exact location of all the genes was obtained from NCBI, http://www.ncbi.nlm.nih.gov in June-July 2005 (build 35.1). A total of 1041 genes were used, and some of the key genes are illustrated in the top lanes of Figure 2–5 and the associated pathways in Figure 1.

### Statistical analyses

Both parametric and non parametric methods were used, in particular Pearson Product Moment Correlation Coefficient (PMCC) *r*, Spearman’s rank correlation coefficient ρ and Kendall’s coefficient of rank correlation for tied ranks ***τ***_b_ were calculated with the help of the statistical software (SPSS 13.0). Significance was tested. Two-tailed p values were reported and the significance was assessed at the 5%, 0.1% and 0.001% level.

PMCC is the ordinary product moment correlation of two variables x and y and it is a measure of how well a linear equation describes the relation between the two variables on the same object. It is defined as the covariance between x and y divided with the product of their standard deviations and can take on values from −1 (perfect inverse correlation) via 0 (no correlation) to +1 (perfect correlation).

When a number of individuals are arranged in order according to some quality, which they all possess to a varying degree, they are said to be ranked. Spearman’s ρ is a product moment correlation coefficient much like Pearson’s *r*. The difference is that Spearman’s ρ can be used when the participants are ranked on both variables.

In practical applications of ranking methods there sometimes arise cases in which two or more objects or individuals are so similar that no preference can be expressed between them. The members are then said to be tied. If the data contain a lot of such ties, other methods are more appropriate to use, such as Kendall’s ***τ***_b_ that uses a correction for this. All three methods are restricted to the interval −1 to +1.

## Results

All the eight data sets are compared on the four characteristic chromosome numbers 3, 8, 14 and 17, that in the present study were here found to be typical for chromosomes 1-3, 4-12, 13-15 and 16-22, as shown in Figs 2–5 respectively. There is often a close agreement between regions with a high chiasma, MLH1 immunofluorescence, and recombination density with sites having a high probability for localized RIBs after X-ray or neutron irradiation as also seen in Fig 6 for all autosomal chromosomes. To get a more accurate quantitative description different numerical statistical measures are applied in the following subsections. Finally some of the key findings are shown for all autosomal chromosomes in Figs 6 and 7

**Figure 6.**
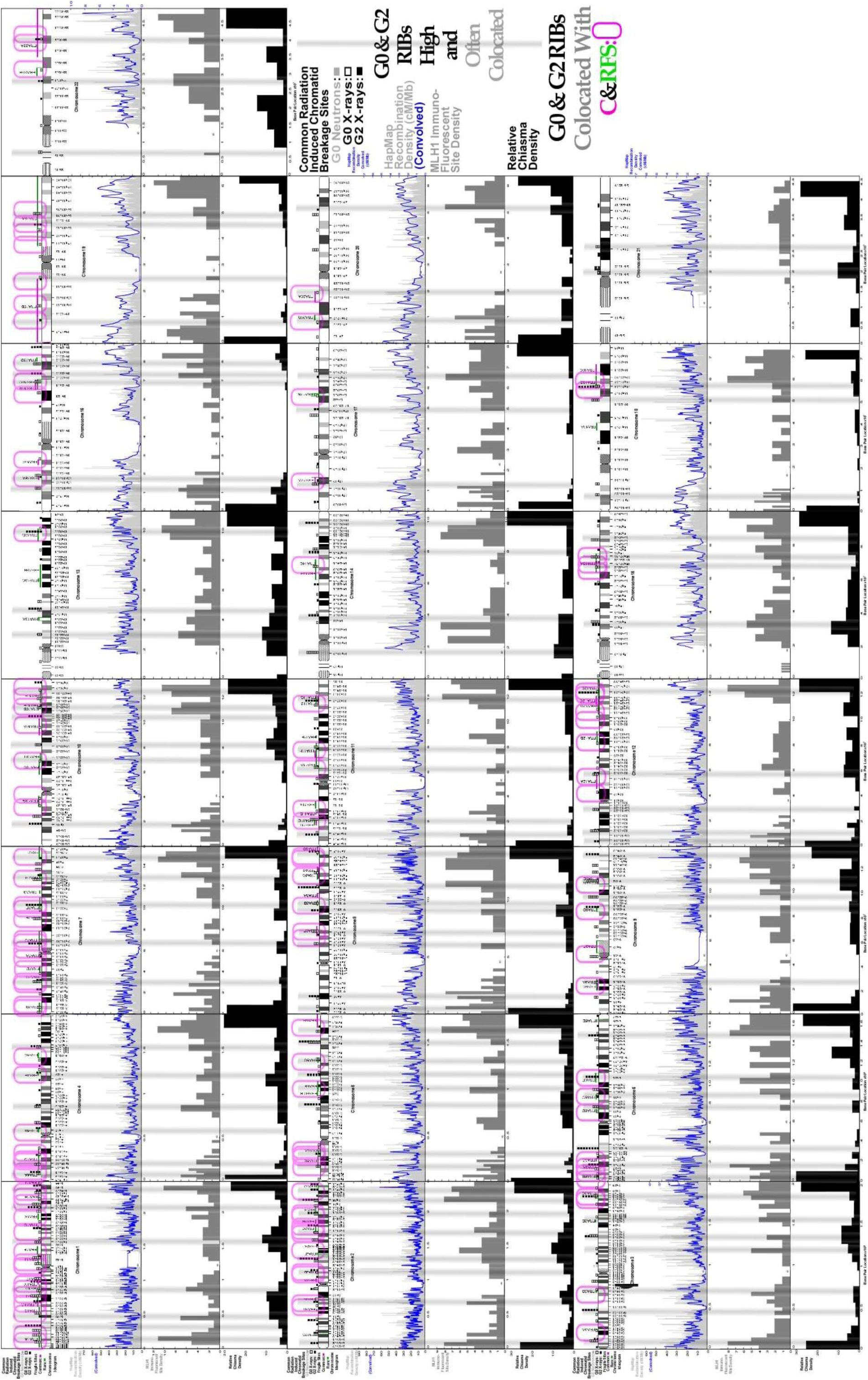
Comparison of common radiation induced chromatid breakpoints and fragile sites with HapMap recombination density, the MLH1 and chiasma distributions for all 22 autosomal chromosomes. Cf. legend of Figure 2–5. It is seen that many common RIBs (≈29%) are located within fragile sites. Furthermore, almost half of all common RIBs (≈49%) are either of rather high intrinsic probability (Gray shading ≈17%) or collocated with common and rare fragile sites (Pink ellipses ≈33%).

**Figure 7.**
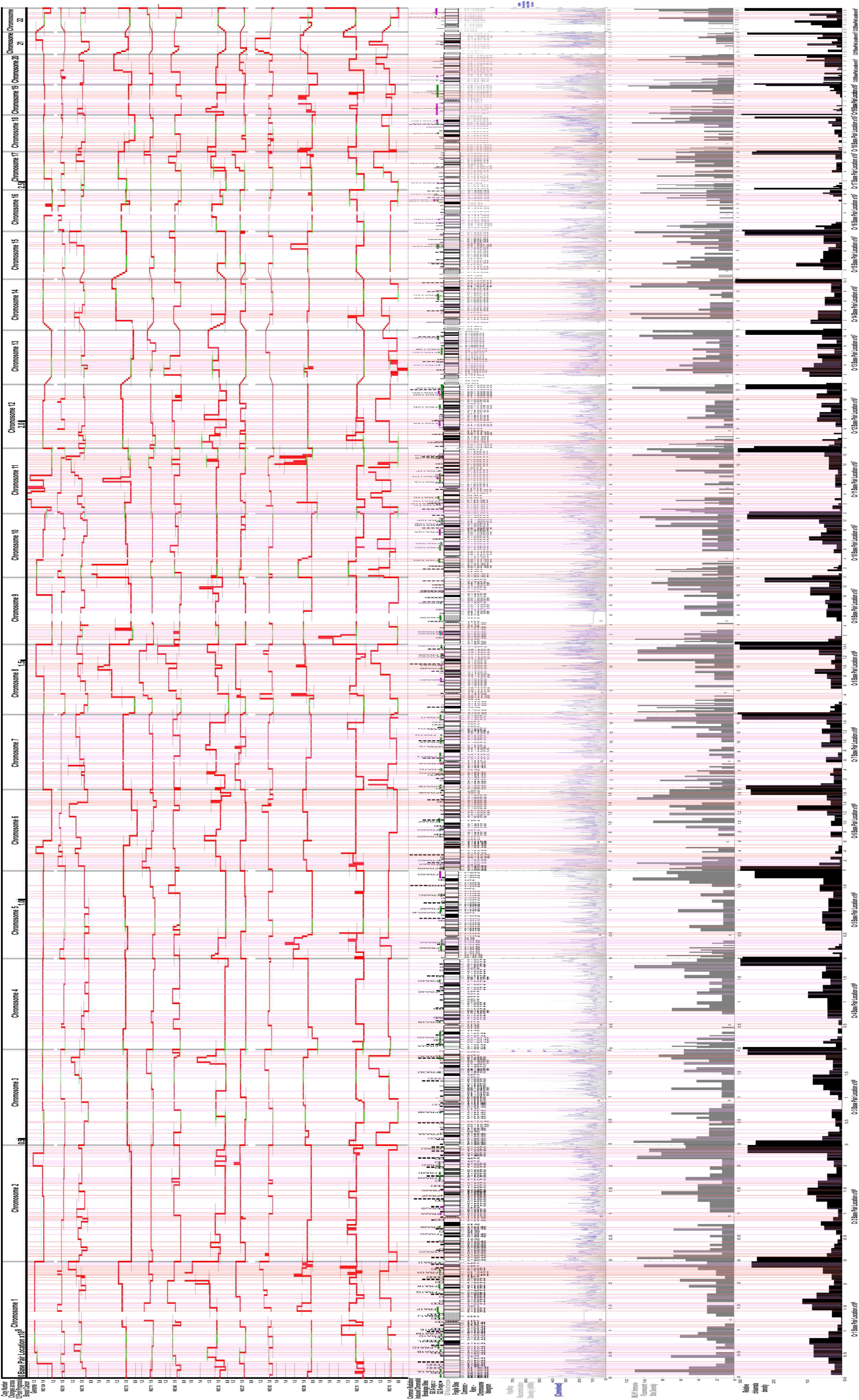
The location of major CNCs (Red histograms) for the whole set of 22 autosomal chromosomes, for 12 poor prognoses complex sawtooth type breast cancer patients, are compared with common RIBs and FSs. It is seen that a large percentage of the CNCs are located precisely at common RIBs or inside the regions of FSs (Pink translucent lines) and often grouped together as denser regions. In some cases, two or more CNC breaks from different patients are collocated over the same RIB or FS and in about 10 % of these cases they are instead located outside these regions (Red translucent lines). In extreme cases, as many as 38% of the CNCs are located at X-ray and neutron RIBs, and 41% on the common and rare FS regions (patient HZ31 second from bottom). Below the chromosome ideograms the HapMap, MLH1 immunofluorescence and Chiasma densities are also shown to indicate the possible association of CNCs with recombination phenomena.

### Correlation between chiasma, MLH1 immunofluorescence, and HapMap recombination sites

There is obviously a strong correlation between chiasma, MLH1 immunofluorescence, and HapMap recombination sites across the genome. The mean correlation coefficient between chiasma locations and MLH1 binding sites was 0.466 including only coefficients that are statistically significant (Bold in Table 1). Even higher coefficients, i.e. showing a stronger correlation, were found when comparing the recombination density distribution from the HapMap project with chiasma sites and MLH1 hot spots, 0.491 and 0.532 respectively. Correlation coefficients and corresponding p values for individual chromosomes are given in Table 1.

**Table 1:**
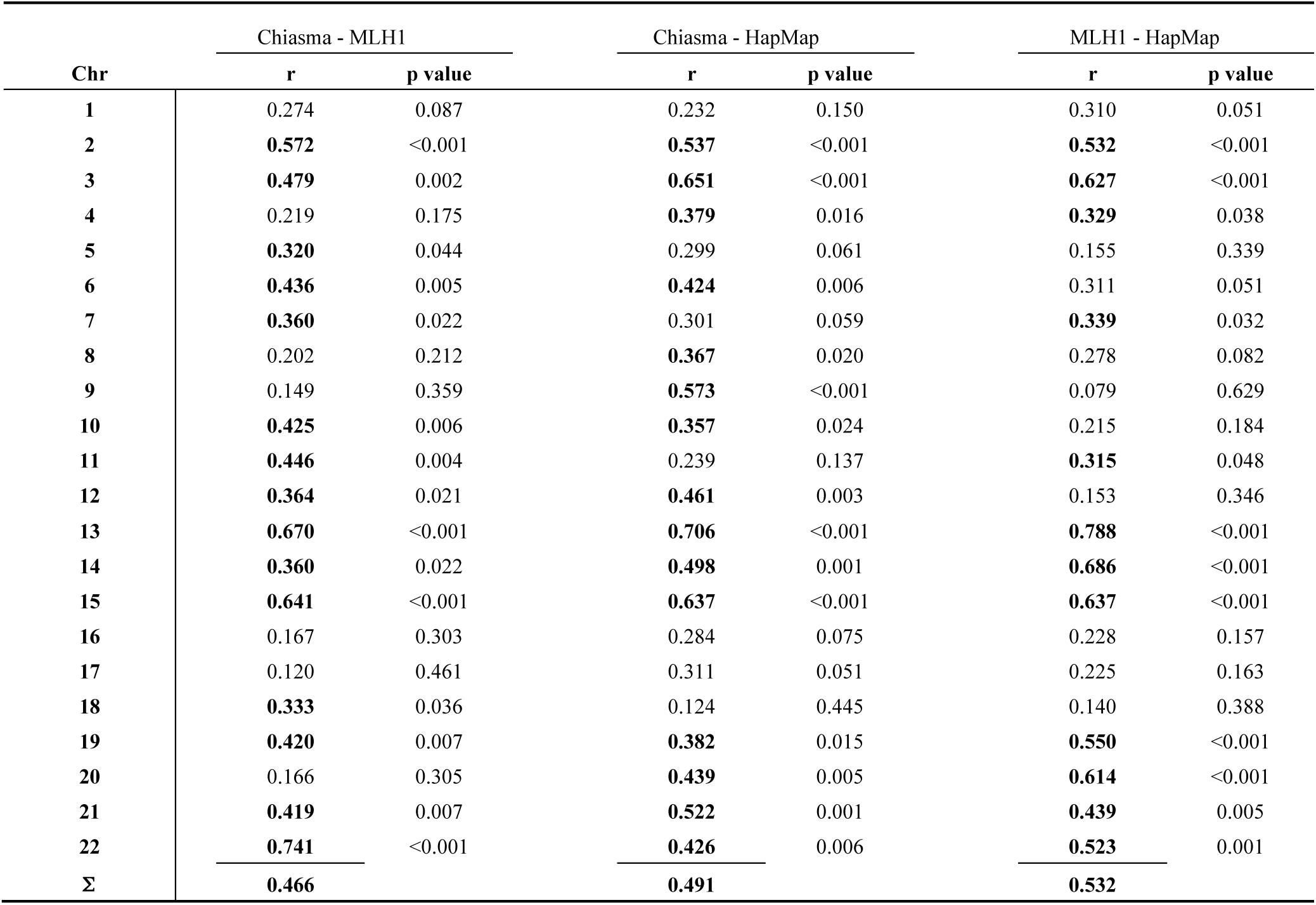
Correlation between chiasma, MLH1 immunofluorescence, and HapMap recombination sites for individual chromosomes. Bold figures are significant with p<0.05.

### Correlation between HapMap recombination density and CNCs in cancer cells

Another interesting and possible relation is the one between the recombination probability and CNC. Because there are many CNCs on chromosome 8 this chromosome was chosen as an example. As can be seen in the left half of Table 2, the ROMA amplitude decreases weakly but significantly with the recombination rates for patients 5, 13 and 16, using Pearson *r* (p<0.05), and for patients 1, 2, 5, 7, 10, 11, 13, 15, 16 and 22, using Kendall’s ***τ***_*b*_ (p<0.001). The same result was obtained with Spearman’s *ρ* as with Kendall’s ***τ***_*b*_, with the exception of patient 11, showing only a very weak positive correlation (p<0.001). In general, the correlation between recombination density from the HapMap project and ROMA amplitude showed a weak negative relationship, i.e. CNC are overrepresented in chromosomal regions with lower than average recombination density.

### Correlation of CpG density with CNCs in cancer cells

The CpG density was calculated at interval lengths of 10 kb on each side of the detected ROMA CNC region. The right half of Table 2 shows the result of this correlation. The table shows a weak, statistically highly significant invers relationship between ROMA amplitude and CpG density for all patients except 12, 13 and 20 using Pearson’s *r* (p<0.001) and for all patients except 12 and 20 using Kendall’s ***τ***_*b*_ (p<0.01) and Spearman’s *ρ* (p<0.01). Patient 20 shows a weak positive relation when using Kendall’s ***τ***_*b*_ and Spearman’s *ρ* (p<0.05). Thus, the ROMA amplitudes are negatively correlated with CpG density in humans, i.e. the ROMA breakpoints show as may be expected a tendency to be located in regions with lower than average CpG density.

### Correlation of HapMap recombination density distribution and cancer related genes

The over representation of CNC in regions with lower than average recombination density may suggest that the mechanisms of crossing-over are most common in regions absent of cancer-related genes. Testing this hypothesis by correlating the recombination density with genes related with cancer proved to be very fruitful. When all 1027 genes were included, a clear majority of the genes were located in regions with a recombination density lower than average. Table 3 is showing this result in the first column. Testing the same hypothesis for different subgroups of genes, such as the oncogenes, the tumor suppressor genes and the DNA repair genes, the result was even more obvious. For the oncogenes only chromosome 14 showed a result that was contradicting, with 4 genes out of 7 located in a region of higher than average recombination density (Table 3).

**Table 3:** The number and percentage of cancer related genes located at lower than average HapMap recombination frequency for each individual chromosome

| Chr | mean | Cancer - Related Genes |  |  | Oncogenes |  |  | Tumor Suppr. Genes |  |  | DNA Repair Genes |  |  |
| --- | --- | --- | --- | --- | --- | --- | --- | --- | --- | --- | --- | --- | --- |
|  |  | total | lower | % | total | lower | % | total | lower | % | total | lower | % |
| 1 | 1.340 | 103 | 82 | 79.6 | 19 | 15 | 78.9 | 18 | 13 | 72.2 | 8 | 8 | 100.0 |
| 2 | 1.160 | 63 | 49 | 77.8 | 9 | 7 | 77.8 | 8 | 5 | 62.5 | 8 | 6 | 75.0 |
| 3 | 1.253 | 76 | 58 | 76.3 | 17 | 12 | 70.6 | 15 | 11 | 73.3 | 10 | 10 | 100.0 |
| 4 | 1.175 | 25 | 20 | 80.0 | 9 | 8 | 88.9 | 2 | 1 | 50.0 | 3 | 3 | 100.0 |
| 5 | 1.220 | 47 | 42 | 89.4 | 4 | 2 | 50.0 | 9 | 8 | 88.9 | 11 | 10 | 90.9 |
| 6 | 1.215 | 52 | 42 | 80.8 | 13 | 11 | 84.6 | 7 | 6 | 85.7 | 6 | 5 | 83.3 |
| 7 | 1.295 | 68 | 62 | 91.2 | 7 | 6 | 85.7 | 11 | 10 | 90.9 | 8 | 5 | 62.5 |
| 8 | 1.325 | 45 | 35 | 77.8 | 12 | 9 | 75.0 | 7 | 4 | 57.1 | 9 | 8 | 88.9 |
| 9 | 1.735 | 41 | 33 | 80.5 | 7 | 4 | 57.1 | 6 | 5 | 83.3 | 5 | 4 | 80.0 |
| 10 | 1.434 | 34 | 29 | 85.3 | 12 | 11 | 91.7 | 8 | 6 | 75.0 | 6 | 5 | 83.3 |
| 11 | 1.277 | 82 | 60 | 73.2 | 15 | 11 | 73.3 | 21 | 15 | 71.4 | 12 | 12 | 100.0 |
| 12 | 1.466 | 56 | 42 | 75.0 | 9 | 6 | 66.7 | 5 | 3 | 60.0 | 8 | 8 | 100.0 |
| 13 | 1.332 | 26 | 22 | 84.6 | 1 | 1 | 100.0 | 10 | 7 | 70.0 | 3 | 3 | 100.0 |
| 14 | 1.404 | 27 | 16 | 59.3 | 7 | 3 | 42.9 | 0 | 0 | 0.0 | 8 | 6 | 75.0 |
| 15 | 1.569 | 33 | 31 | 93.9 | 2 | 2 | 100.0 | 7 | 6 | 85.7 | 5 | 4 | 80.0 |
| 16 | 1.743 | 30 | 17 | 56.7 | 2 | 2 | 100.0 | 5 | 2 | 40.0 | 4 | 2 | 50.0 |
| 17 | 1.688 | 75 | 59 | 78.7 | 9 | 6 | 66.7 | 8 | 6 | 75.0 | 11 | 9 | 81.8 |
| 18 | 1.573 | 15 | 15 | 100.0 | 2 | 2 | 100.0 | 4 | 4 | 100.0 | 1 | 1 | 100.0 |
| 19 | 1.886 | 58 | 40 | 69.0 | 14 | 9 | 64.3 | 5 | 3 | 60.0 | 9 | 9 | 100.0 |
| 20 | 1.860 | 40 | 31 | 77.5 | 10 | 6 | 60.0 | 2 | 2 | 100.0 | 2 | 2 | 100.0 |
| 21 | 1.927 | 5 | 4 | 80.0 | 2 | 2 | 100.0 | 1 | 0 | 0.0 | 0 | 0 | 0.0 |
| 22 | 2.327 | 26 | 25 | 96.2 | 3 | 3 | 100.0 | 8 | 8 | 100.0 | 3 | 3 | 100.0 |

Almost the same result was obtained for the Tumor Suppressor Genes. On all autosomal chromosomes except chromosomes 16, 21 and possibly 14, the majority of the genes were situated in areas with a decreased recombination rate (Table 3). The most fascinating result was obtained for genes involved in DNA repair. In this study 123 out of a total of 140 genes, i.e. 88%, were located in DNA regions with no or lower than average recombination density (Table 3).

### Correlation of CpG density and cancer related genes

It is well known that a high CpG density is often associated with a high gene density in the same region. But does that hold also for a certain group of genes, for example the oncogenes? By investigating the correlation between the CpG densities around 1026 cancer related-genes, the result in Table 4 were obtained. It can be seen from Table 4 that a higher than average CpG density (almost 83%) is located around cancer related genes in agreement with the earlier results. By studying the same relation, but only for oncogenes, an even higher CpG density (almost 86%) was found on average (Table 4). With a chromosome mean CpG density ranging from 0.0076 to 0.0161 or around 1%, 159 out of 185 genes, i.e. more than 85%, were located in regions of higher than average CpG density. Similar results were obtained for the tumor suppressor genes and the DNA repair genes (Table 4).

**Table 4:** The number and percentage of cancer related genes located in higher than average CpG density for each individual chromosome at a CpG Window of 2500 kbp.

| Chr | mean | Cancer-Related Genes |  |  | Oncogenes |  |  | Tumor Suppr. Genes |  |  | DNA Repair Genes |  |  |
| --- | --- | --- | --- | --- | --- | --- | --- | --- | --- | --- | --- | --- | --- |
|  |  | total | higher | % | total | higher | % | total | higher | % | total | higher | % |
| 1 | 0.0091 | 103 | 83 | 80.6 | 19 | 14 | 73.7 | 18 | 13 | 72.2 | 8 | 7 | 87.5 |
| 2 | 0.0087 | 63 | 51 | 81.0 | 9 | 8 | 88.9 | 8 | 7 | 87.5 | 8 | 7 | 87.5 |
| 3 | 0.0080 | 76 | 65 | 85.5 | 17 | 13 | 76.5 | 15 | 13 | 86.7 | 10 | 10 | 100.0 |
| 4 | 0.0076 | 25 | 20 | 80.0 | 9 | 8 | 88.9 | 2 | 2 | 100.0 | 3 | 3 | 100.0 |
| 5 | 0.0081 | 47 | 32 | 68.1 | 4 | 3 | 75.0 | 9 | 6 | 67.7 | 11 | 7 | 63.6 |
| 6 | 0.0084 | 52 | 44 | 84.6 | 13 | 10 | 76.9 | 7 | 6 | 85.7 | 6 | 6 | 100.0 |
| 7 | 0.0098 | 68 | 45 | 66.2 | 7 | 4 | 57.1 | 11 | 6 | 54.5 | 8 | 6 | 75.0 |
| 8 | 0.0087 | 45 | 35 | 77.8 | 12 | 10 | 83.3 | 7 | 5 | 71.4 | 9 | 8 | 88.9 |
| 9 | 0.0085 | 40 | 36 | 90.0 | 7 | 6 | 85.7 | 6 | 5 | 83.3 | 5 | 5 | 100.0 |
| 10 | 0.0099 | 34 | 28 | 82.4 | 12 | 11 | 91.7 | 8 | 6 | 75.0 | 6 | 5 | 83.3 |
| 11 | 0.0095 | 82 | 67 | 81.7 | 15 | 14 | 93.3 | 21 | 15 | 71.4 | 12 | 10 | 83.3 |
| 12 | 0.0096 | 56 | 47 | 83.9 | 9 | 8 | 88.9 | 5 | 4 | 80.0 | 8 | 7 | 87.5 |
| 13 | 0.0069 | 26 | 26 | 100.0 | 1 | 1 | 100.0 | 10 | 10 | 100.0 | 3 | 3 | 100.0 |
| 14 | 0.0079 | 27 | 26 | 96.3 | 7 | 7 | 100.0 | 0 | 0 | 0 | 8 | 7 | 87.5 |
| 15 | 0.0086 | 33 | 31 | 93.9 | 2 | 2 | 100.0 | 7 | 7 | 100.0 | 5 | 5 | 100.0 |
| 16 | 0.0121 | 30 | 25 | 83.3 | 2 | 2 | 100.0 | 5 | 4 | 80.0 | 4 | 4 | 100.0 |
| 17 | 0.0146 | 75 | 54 | 72.0 | 9 | 8 | 88.9 | 8 | 7 | 87.5 | 11 | 6 | 54.5 |
| 18 | 0.0088 | 15 | 12 | 80.0 | 2 | 2 | 100.0 | 4 | 3 | 75.0 | 1 | 1 | 100.0 |
| 19 | 0.0161 | 58 | 57 | 98.3 | 14 | 14 | 100.0 | 5 | 4 | 80.0 | 9 | 9 | 100.0 |
| 20 | 0.0113 | 40 | 36 | 90.0 | 10 | 9 | 90.0 | 2 | 2 | 100.0 | 2 | 2 | 100.0 |
| 21 | 0.0078 | 5 | 5 | 100.0 | 2 | 2 | 100.0 | 1 | 1 | 100.0 | 0 | 0 | 0 |
| 22 | 0.0112 | 26 | 26 | 100.0 | 3 | 3 | 100.0 | 8 | 8 | 100.0 | 3 | 3 | 100.0 |
| **Σ** |  | **1026** | **851** | **82.9** | **185** | **159** | **85.9** | **167** | **134** | **80.2** | **140** | **121** | **86.4** |

### Correlation between RIBs and FSs

As shown in Fig 6 for the whole set of 22 autosomal chromosomes the X-ray and neutron induced breakpoints are commonly located in the regions of common and rare FS. In fact about 72 & 45 out of 205 & 156 or about 36&30% of the common G0 & G2 RIBs are collocated with 54 & 45 out of 120 or about 45 & 37% of the FSs respectively. Furthermore about 50% of the RIBs are either collocated or highly probable and located in regions of high chiasma MLH1 immunofluorescence, and HapMap recombination density as seen in Fig 6.

### Correlation between RIBs and FSs with CNCs in Severely Malignant Breast Cancer

The CNC distribution of twelve breast cancer patients with very poor prognosis^23^ (complex type I or sawtooth), are found to be highly correlated with X-ray and neutron RIBs as well as common and rare FS, as shown in Fig 7 for the whole set of 22 autosomal chromosomes. For the patient with most CNCs namely WZ31 more than 75% of the CNCs are located in these regions. In average over the 12 patients 69% of the CNCs are located at X-ray RIBs and 49% of the CNCs are in the common and rare FS regions. This indicates that about 18% of the CNCs are simultaneously located at RIBs and FS. In addition, about 20% of the CNCs are collocated with CNCs of the other patients and at one site the CNCs are collocated for eight patients. Only 3% of the CNCs are alone and the mean multiplicity at colocation is 3.2.

## Discussion

The present study support that there are many different regions in the genome with an increased probability of changes and rearrangements. Eight different affected data sets covering the whole human genome have been compared with regard to their genomic distribution and a close correlation between these have been found for all autosomal chromosomes.

### Chiasma, MLH1 immunofluorescence, and HapMap recombination sites

The three lower lanes two of which are histogram like in Figure 2-6 with considerably lower resolution than the HapMap set shown also convolved with a Gaussian filter to more clearly show gross structures. It is seen that the three datasets have a large degree of variability partly associated with the technical development over the last 40 years and the differences between male and female data, even though they show many common features. Some of the deviations may be due to the fact that HapMap data generally are averages over a large number of males and females^18–21^ whereas MLH1 and chiasma distributions generally pertain to males^3, 26^. This is seen particularly in the first three Figs 2-4 for the chromosomes 3, 8 and 14. This is often associated with a high CpG density indicating that this is not a sufficient property for avoiding local alterations and breaks. This indicates that the larger HapMap data set may be more generally applicable since it averages over a larger number of patients of both sexes so individual variations reduce the correlation between the smaller male MLH1 and chiasma data sets. The key aim of the formation and the subsequent repair of massive DSBs^27^ is the generation of a few meiotic crossover events along each chromosome. The determinant for the crossover localization is the distribution of the DSB that initiate the recombination. In mammals, as well as in other organisms it is highly likely that DSB also initiate meiotic recombination. The central protein in DSB formation, Spo11 is highly conserved^28^. Meiotic cells make an excess of DSB compared to the number of recombination events observed in other cells. Available data indicate that the number of non-crossover events is 10 times higher than the crossover events^29^. While the distribution of the recombination sites is clearly nonrandom, the preference for certain locations might be associated with the significance and function of different regulatory regions and groups of genes in the regions involved. When there is a mechanism selecting where the breaks are induced, it is likely that regions that are more tolerant to alterations host the recombination events at the same time as the mismatch repair machinery may be linked to these sites.

When a meiotic cell induces many hundreds of breaks on its own genome, and a single unrepaired break may be lethal, high fidelity DSB repair is a critical process for high fidelity cell survival. This may explain why MLH1 mismatch repair gene-binding sites are incorporated at the crossover events. A strong correlation was confirmed between the MLH1 foci and cytological detected chiasma sites (Pearson correlation coefficient = 0.466; p<0.05) as well as with the recombination probability distribution from the HapMap project (Pearson correlation coefficient = 0.532; p<0.05; Table 1). Besides these large-scale tendencies it is also seen that there are a fair amount of local deviations, probably indicating that there may be other sources of individual variability in the radiation induced breakage distributions as well as the MLH1 and chiasma distributions. This is also seen in the recent MLH1 data sets^26^ and the larger variability seen in the HapMap data even when convolved with a Gaussian filter to emphasize gross structure. To address the relation between recombination and cancer associated genes, we have calculated the recombination rate at the gene locations, for all autosomal chromosomes. The recombination rate is on the average lower in the genes, with about 80% (814 of 1027) being located in regions of lower than average recombination probability. This is also supported by the result from Myers et al^8^, who investigated the relation between recombination and genes in general. The recombination hotspots in humans seem to occur near genes but are preferentially located outside the transcribed domain^8^.

### RIBs and FSs

It is well known that the initial DNA damage caused by radiation is essentially randomly distributed across the whole genome at least for low ionization density electrons and photons, whereas chromosomal breakpoints shown in Figs 2-7 are generally not. This indicates that these locations somehow are more prone to develop chromosomal breaks during the DNA repair process based on an initial largely random DSB damage site distribution. Not surprising, some of the FS are covering the RIBs as seen in two lanes above chromatid breaks in Figs 2–5 and not least in Fig 6 for all autosomal chromosomes. This indicates that at least some of the RIBs are some kind of hot spots inside the common and rare FSs. Interestingly, Table 3 shows that the DNA repair genes are predominantly avoiding regions with a high recombination rate. This indicates that the repair genes are of critical importance for the survival of the cell when damaged, and for this reason are commonly located areas with a low local recombination probability. This is probably a result of natural selection, since if they were affected by erroneous recombination events they may not function after such an event and the cell may not be able to repair damage and it will often be lost from further divisions. As has been shown recently, intact NHEJ and HR pathways can repair as many as 99% or more of the radiation induced simple DSB^30–32^ and probably even more of SSB and milder damage types, so it is clearly very important for cellular survival that these repair genes are well functioning. Interestingly, the RIB data^9–10^ indicates that at most 1.6 and 3.2 % of the DSBs may result in chromosome breaks during G0 and G2 irradiation. The increase is partly expected due to higher sensitivity during the late G2 phase. This repair efficiency applies to common doses and dose rates in radiation therapy of around 2 Gy but is significantly reduced at very high doses and dose rates when NHEJ have problem to correctly combine the associated broken strands due to a high density of local damage. It is also interesting to note that NHEJ together with Non-allelic homologous recombination (NAHR), Fork Stalling and Template Switching (FoSTeS), Break Induced Replication (BIR) and Microhomology-Mediated BIR (MMBIR) belong to the main mechanisms for human CNCs, genomic instability and rearrangements^33^. Several independent studies have shown that RIBs are preferentially located in R-bands and light G-bands. These are chromosome regions with more open chromatin that are generally gene-rich with high CpG dencity. If the damage is induced in the same location as a recombination hot spot, the likelihood for the break to be repaired is increased. The production of Dual DSBs^30^ and the formation of inter-chromosomal crosslinks, due to misrepair of DSBs, are major causes of death for cells exposed to ionizing radiation. The distribution of recombination hot spots may therefore have an impact on the chromosome aberration formation spectra and the survival of cells exposed to ionizing radiation. We found that many of the RIBs coincide with recombination hot spots as well as with an elevated CpG density, and in many cases also with FS. Due to the limitations in the exact location of breakpoints, and because the FS are generally only mapped to a cytological band of a chromosome, a more detailed correlation analysis of such data would be highly desirable in the future, but more complicated to perform.

### CpG density

To investigate whether copy number variation in cancer cells are associated with hotspots, we calculated the recombination rate at the microarray-detected breaks. In general, there was a weak negative relationship. These gene amplifications and deletions seem overrepresented in chromosomal regions with a lower recombination rate than the chromosome average. It is well known that chromosomal regions of high CpG density exhibit higher gene densities. These regions also contain a higher density of CpG islands. CpG islands influences weather adjacent genes should be active or not depending on the degree of methylation of the surrounding CpG regions. One would therefore expect cancer gene amplifications to be negatively correlated to the CpG density. A highly significant weak negative correlation was found. This is no surprise since a high CpG density is often linked to regions of the genome that are well preserved as discussed above^5, 15, 16^. The ROMA breakpoints thus showed a tendency to be located in regions with lower than average CpG density unlike normal genes and in particular unlike the DNA repair genes. As seen in Table 3, the DNA repair genes have the highest probability to be located in higher than average CpG rich regions. As expected there is a highly increased CpG density for chromosome localizations around oncogenes and DNA repair genes but also around other cancer related genes.

### The Relation between CNCs in poor prognosis Breast Cancers and RIBs and FS

It is not surprising that many of the CNCs are located inside the regions of the FSs. However, the location of even more of the CNCs fall exactly at RIBs may be more unexpected and a new finding. This indicates that the RIBs are really locations on genome that are extra sensitive to damage and more difficult to repair! This may also be concluded from the fact that several of the RIBs are also located within the regions of the FS and may be some kind FS local hot spots. This group of cancer patents is generally receiving radiation therapy. However, the tumor surgery is generally taking place before the radiation treatment, so the patients CNCs should not be affected by later high therapeutic doses. It is of course possible that the natural background radiation and diagnostic imaging like mammography, may have been the reason for some of the CNCs at RIBs. At least the former source is rather unlikely, but a 2 mGy of mammography dose will induce about 0.1 DSBs per cell or about one DSB in every 10 cells of the imaged volume, which may have had some effect after many years of repeated diagnostics. Premature chromosome condensation may be a method to study genetic tissue changes after longer periods of tissue exposer.

### Recent Related Studies

The recently published paper on the mutational heterogeneity in cancer^34^ observed similar variations across the genome in mutation probability as we see in DNA break induction and recombination probability. Furthermore, they also found that it varies widely over the genome and correlates well with DNA replication time and expression level (cf their Figs 3a and S3 with Figure 4 above). The recent study of aphidicolin (a DNA polymerase inhibitor) induced fragile sites^35^ and genetic divergence across the human genome^36^ seams to largely show similar correlations as we see in the present study. Furthermore, the genome-wide study of hot spots of DNA double-strand breaks in human cells^37^ not surprisingly show a strong correlation not least with the presently studied radiation-induced breaks, the CpG density and the Hap Map data. Similarly, the recent study of breakage in specific chromosome bands after acute exposure to oil and benzene show similar correlations and close association to fragile sites^38^. The recent genome wide study of p53 pathway cancer related Single Nucleotide Polymorphisms and expression quantitative trait loci^39^ both agree very well with the presently described radiation-induced Chromatid Breakage points and both types of Fragile Sites. This may also be expected due to the genome wide influence of different types of increased local genomic instability. The key gens FHIT (genome “caretaker” located at FRA3B, see Fig 1), WWOX (involved in DNA damage checkpoint activation, located at FRA16D) and p53 (genome “guardian”) are frequently involved in commonly activated fragile sites and early events in cancer induction^40^ as well as in DNA damage repair^41^. In general, the fragile sites often harbor tumor suppressor genes that are active in the DNA damage response pathways^42^.

## Conclusions

It is compelling that common RIBs, CNCs, chiasma and FSs^43^ are all strongly dependent on how the cells are handling DSB and more generally DNA damage on their genome. From this point of view, looking at the whole genome (Figs 6, 7) it is striking how often the common RIBs fall inside the fragile sites and exactly at CNCs loci. In average 33% of the common RIBs are collocated with 41% of the FSs respectively, and then practically always in regions of elevated recombination, MLH1 immunofluorescence and chiasma density as seen in Fig 6. The observed large-scale correlation between HapMap recombination sites, chiasma, MLH1 immunofluorescence and highly probable RIBs as seen in Figures 2-6 is striking, even though there seems too be some local deviations. This may also be expected due to use of different population averages in the former datasets, such as when using germ cells from males or females and cells from other tissues, but also due to the wide range of genomic variability over the human genome and the associated genomic instability and hyper mutation in human cancers^32–44^. In breast cancer patients with poor prognosis^23^ on average 69% of the CNCs are located at X-ray RIBs covering less than 20% of the genome, whereas 49% of the CNCs are located in the common and rare FS regions covering about 27% of the genome. The RIBs are thus more localized and about 2 times more likely per Mbp to cause CNC then the FS regions, but otherwise have similar properties as they are linked to an increased risk of local breaks and rearrangements of the genome. This may be due to a local DSB repair problem with increased risk for miss repair. The present study has also shown that only 3% of 782 CNCs are alone outside these regions (cf Fig7) and missing other CNCs, in severely malignant breast cancers. Many of the RIBs may therefore be regarded as a new type of more localized DNA repair fragility likely to be involved also in other types of cancers, and most likely, without the need for exposer to ionizing radiations^9–12^ or benzene^38^ that first identified their existence.

